# Landscape of gene transposition-duplication within the Brassicaceae family

**DOI:** 10.1101/236299

**Authors:** Dong-Ha Oh, Maheshi Dassanayake

## Abstract

We developed the CLfinder-OrthNet pipeline that detects co-linearity in gene arrangement among multiple closely related genomes; find ortholog groups; and encodes the evolutionary history of each ortholog group into a representative network (OrthNet). Using a search based on network topology, out of a total of 17,432 OrthNets in six Brassicaceae genomes, we identified 1,394 that included gene transposition-duplication (*tr-d*) events in one or more genomes. Occurrences of *tr-d* shared by subsets of Brassicaceae genomes mirrored the divergence times between the genomes and their repeat contents. The majority of *tr-d* events resulted in truncated open reading frames (ORFs) in the duplicated loci. However, the duplicates with complete ORFs were significantly more frequent than expected from random events. They also had a higher chance of being expressed and derived from older *tr-d* events. We also found an enrichment, compared to random chance, of *tr-d* events with complete loss of intergenic sequence conservation between the original and duplicated loci. Finally, we identified *tr-d* events uniquely found in two extremophytes among the six Brassicaceae genomes, including *tr-d* of *SALT TOLERANCE 32* and *ZINC TRANSPORTER 3.* The CLfinder-OrthNet pipeline provides a flexible and a modular toolkit to compare gene order, encode and visualize evolutionary paths among orthologs as networks, and identify all gene loci that share the same evolutionary history using network topology searches.

**Funding source:** This work was supported by National Science Foundation (MCB 1616827) and the Next Generation BioGreen21 Program (PJ011379) of the Rural Development Administration, Republic of Korea.

**Online-only Supplementary materials** includes supplementary text (S1-S10), methods (M1-M4), figures (S1-S7), and tables (S1-S3), in two PDF files, one for text and methods and the other for figures and tables. Additionally, Supplementary Dataset S1 is available at the Figshare repository (https://doi.org/10.6084/m9.figshare.5825937) and Dataset S2 and S3 as separate Excel files.

## INTRODUCTION

Co-linearity among closely related genomes erodes over time due to the accumulation of mutations including gene duplication, deletion, and transposition (Wicker et al. 2010). Gene duplication affects gene dosage, which may lead to divergence of expression and functions among duplicates (Wang, Wang, et al. 2012; Assis and Bachtrog 2013). Gene transposition events modify expression strength and tissue-specificity through changes in regulatory sequences (Oh et al. 2014; Arsovski et al. 2015), local epigenetic environment (Durand et al. 2012), and proximity to enhancers and chromatin structural contexts (Feuerborn and Cook 2015; Zhu et al. 2015; Hu and Tee 2017). Such events have been associated with variation in copy numbers of genes and transcripts (Oh et al. 2010; Oh et al. 2014), as well as localization (Liu et al. 2014) and functions (Panchy et al. 2016; Wang et al. 2016) of encoded proteins among orthologous genes. A large number of studies have reported examples of gene level duplications and transpositions as key underlying sources for adaptations to specific environments or speciation (Boore et al. 1998; Cook et al. 2012; Kondrashov 2012; Grandaubert et al. 2014; Simon et al. 2015; Gan et al. 2016; Shirai et al. 2017). *De novo* assembled genomes released at unprecedented rates today (Wang et al. 2014; Koenig and Weigel 2015; Du et al. 2017; Wang et al. 2017) enable us to analyze gene gain and loss as well as duplication and transposition among closely related taxa. These resources call for novel methods for systematic comparative analysis of genomes.

Comparative analysis of co-linearity enable identifying modes of gene duplications (Freeling 2009; Emms et al. 2016) and tracing the origin of genes or gene families and their evolutionary history (Vlad et al. 2014; Zhao et al. 2017; Zhao and Schranz 2018). A number of tools are available for identification of gene blocks or large genomic regions co-linear among multiple genomes (Proost et al. 2012; Wang, Tang, et al. 2012; Tang et al. 2015). Another set of tools can be used to identify orthologs in related genomes for a gene of interest and visualize synteny and evolutionary events associated with them (Chen et al. 2006; Lyons and Freeling 2008; Wall et al. 2008; Grin and Linke 2011; Vandepoele 2017). However, currently we do not have a method that can retrieve all ortholog loci within multiple genomes that have likely undergone the same set of evolutionary events without a prior assignment of a gene of interest. To address this, we introduce the CLfinder-OrthNet pipeline, which identifies co-linearity (CL) in the gene order among multiple genomes, identify “ortholog groups” based on co-linearity, and encodes genes in each ortholog group as a network of orthologs (OrthNet). Each ortholog group includes orthologs and paralogs likely derived from a single ancestral locus in multiple target genomes. All evolutionary events in each ortholog group, such as gene duplication, deletion, transposition, and any combination of them, in addition to gene synteny conservation, are captured as the topology of an OrthNet. Our pipeline enables detection of all ortholog groups from multiple genome that seemingly underwent the same evolutionary events, by searching OrthNets essentially based on their topologies.

As a proof-of-concept, we applied the CLfinder-OrthNet pipeline to six Brassicaceae genomes that share the same whole genome duplication history (Haudry et al. 2013), including the model species *Arabidopsis thaliana* (Cheng et al. 2016) and two extremophytes *Schrenkiella parvula* (Dassanayake et al. 2011) and *Eutrema salsugineum* (Wu et al. 2012; Yang et al. 2013). *S. parvula* and *E. salsugineum*, the two most salt-tolerant Brassicaceae species so far tested (Orsini et al. 2010), are biogeographicaly seperated and represent taxa adapted to multi-ion salt strsses in soils near a hypersaline lake in central Anatolia (Oh et al. 2014) and combined salt and freezing stresses in salt pans of high latitude regions in the northern hemisphere (Inan et al. 2004; Amtmann 2009), respectively. These two extremophytes provide optimal models for comparative analyses to study plant adaptations to environmental challenges (Dittami and Tonon 2012; Oh et al. 2012).

The CLfinder-OrthNet pipeline detects any combination of gene synteny conservation, duplications, deletions, and transpositions. For the present study, we focused on the relatively under-studied transposition-duplication (*tr-d*) events (Freeling 2009; Wang et al. 2013), which result in variations in both copy numbers and co-linearity, within the six Brassicaceae genomes. Our pipeline identified *tr-d* events unique to a genome or shared by any subset of the six Brassicaceae genomes, as well as the original donor and duplicate loci in each *tr-d* event including loci with truncated coding regions. Using this pipeline, we aim to identify the landscape of lineage-specific and shared *tr-d* events among the target genomes; test whether there is a signature of selective retention among lineage-specific *tr-d* events; and characterize *tr-d* events unique to the extremophyte genomes, which may have contributed to their adaptive evolution.

## RESULTS

### Patterns of co-linearity erosion within the six Brassicaceae genomes

We selected a set of six genomes with the same whole genome duplication history sampled from the Brassicaceae Lineages I and II for the current study (Figure 1). This set includes the model plant *A. thaliana* (Ath) and its relatives in Lineage I, *A. lyrata* (Aly) and *Capsella rubella* (Cru), as well as *Sisymbrium irio* (Sir) and the two extremophytes, *E. salsugineum* (Esa) and *S. parvula* (Spa) in Lineage II. Fig. 1 shows the phylogenetic relationships of the target species with other published genomes in Brassicaceae based on amino acid sequence alignments of 14,614 homolog clusters (See Methods).

**Figure 1.**
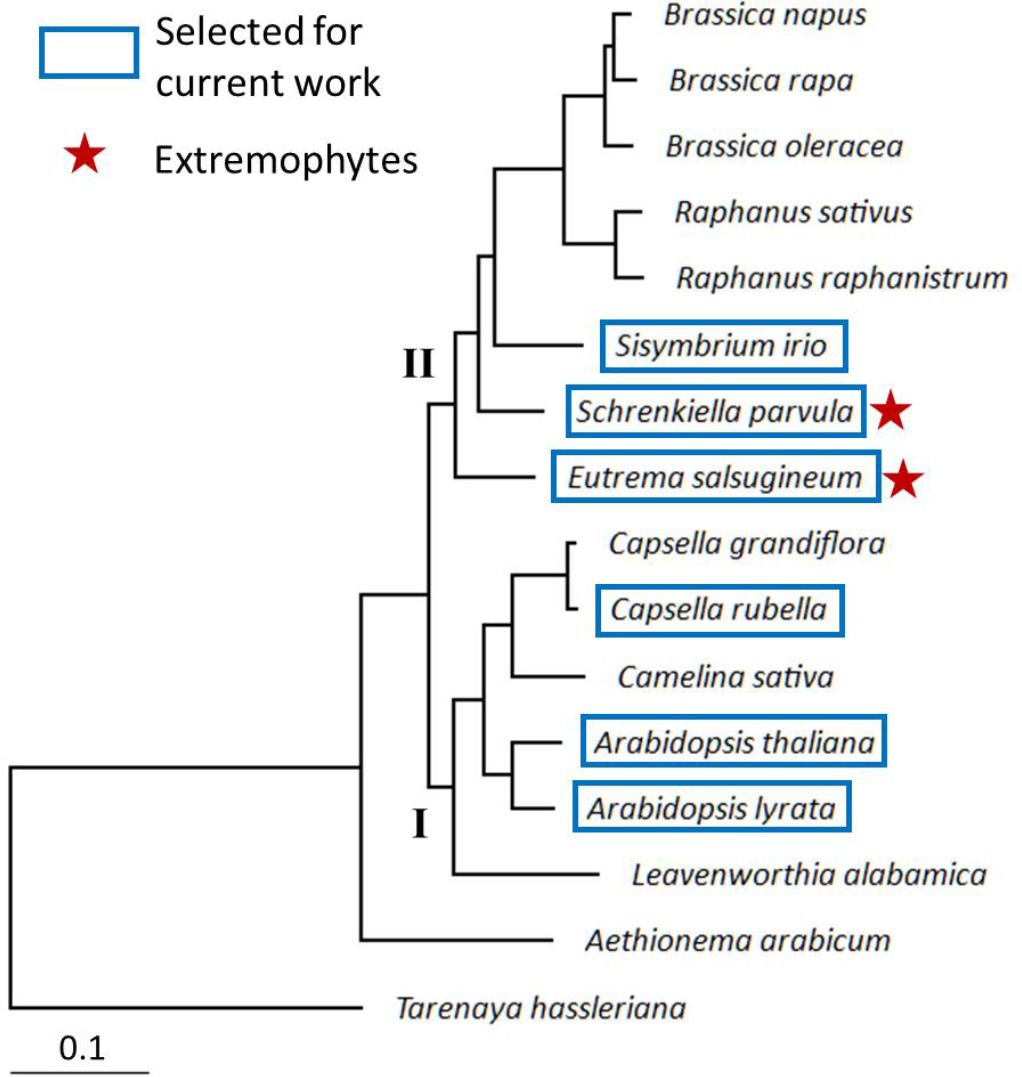
A comparative genomics framework including the two extremophyte/halophyte crucifers, *S. parvula* and *E. salsugineum.* Boxes and stars indicate the six Brassicaceae species selected for this work and halophytes, respectively. The tree was based on an alignment of 14,614 homologous gene clusters, as described in Methods.

Before applying the CLfinder-OrthNet pipeline, we analyzed the degree of co-linearity erosion among the target Brassicaceae genomes by comparing gene orders, as detailed in Supplementary text S1 and Figure S1. Our analysis revealed that two thirds of genes identified as non-transposable element (non-TE) and non-lineage-specific (non-LS) genes in the Brassicaceae genomes showed a conservation of gene order with their immediate neighbors when compared to the genome of *A. thaliana* (Fig. S1C, d_n,n+1_ ≤ 1). The proportion of non-TE and non-LS gene loci showing a proximal (Fig. S1C, dn,n+1 = 2^∼^20) and distal (Fig. S1C, d_n,n+1_ >20 and “Diff Chr”) gene order displacement, compared to their immediate neighbors, was correlated with the divergence time between genomes and their TE contents, respectively (Fig. S1D). This suggested two different mechanisms for co-linearity erosion. The proximal gene order displacements were likely resulted from mutations accumulated over time after the divergence of genomes, while the distal gene order displacements may have been initiated by the presence of repetitive sequences and TE activities (See Supplementary Text S1 and Supplementary Methods M1 for more detail).

### Development of the CLfinder-OrthNet pipeline

Our pipeline consists of two modules: CLfinder and OrthNet (Figures 2 and S2). The first module, CLfinder, compares all possible pairs of query and target genomes and test whether each homologous gene pair (i.e. “best-hit” pair, Supplementary Text Glossary) is co-linear (Figures S2 and S3, and Dataset S1). CLfinder accepts three inputs: representative gene models for all loci in each genome, clusters of paralogs within each genome, and lists of best-hits between all possible query-target genome pairs (Fig. 2 and S2). Users can select the methods and criteria for defining paralog clusters and best-hit pairs, as well as the sensitivity and stringency for the co-linearity detection by controlling three parameters: window_size (*W*), num_CL_trshld (*N*), and gap_CL_trshld (*G*). Based on these parameters, the CLfinder module searches both up- and downstream of each locus in the query genome for “loci-in-chain” based on the order of their best-hits in the target genome, to determine whether a query-target best-hit pair is either co-linear (*cl*), transposed (*tr*), or not able to determine (*nd*) due to the query genome assembly scaffold being too short. When co-linearity was detected only towards one direction, the query-target best-hit pair is considered representing an end of a co-linear genome segment (*cl_end*) derived from inversions, indels, and segmental duplications involving multiple gene loci. A query locus without a best-hit in the target genome is marked lineage-specific (*ls*) (Fig. S3 and Supplementary Methods M2).

**Figure 2.**
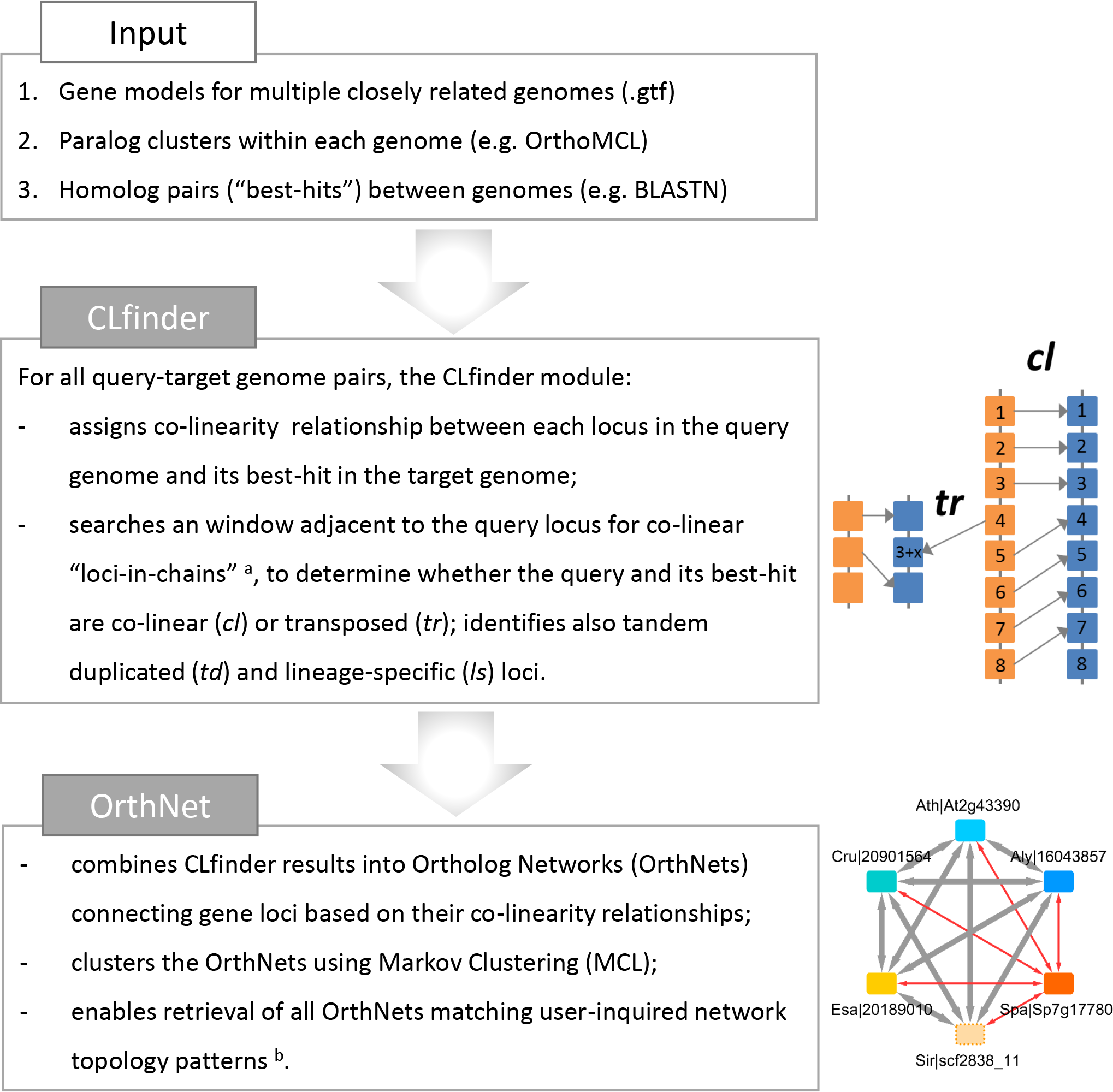
The CLfinder-OrthNet pipeline. ^a^ For the detailed method to determine co-linearity (CL) relationship between the query loci and their most homologous counterpart (“best-hits”) in the target genome, see Figures S2 and S3 and Supplementary Text and Methods. CLfinder output for the six crucifer species highlighted in Fig. 1 are summarized in Table 1, with the full results given as Supplementary Dataset S1. ^b^ See Figures 3-6, for examples of OrthNets with different evolutionary histories represented as different network topologies, e.g. transposition (*tr*) and transposition-duplication (*tr-d*) unique to each species or a group of species.

The second module, OrthNet, combines all pairwise comparisons by CLfinder and encodes co-linearity relationships among orthologs into networks (OrthNets), with gene loci as nodes connected by an edge to their best-hits in other genomes (Figures 2 and 3). Each edge has a property of either co-linear (*cl*), transposed (*tr*), or not determined (*nd*). The *cl* and *tr* edges can be either reciprocal or unidirectional (Fig. 3A, “rc” and “uni”, respectively). OrthNets also include tandem duplicated (*td*) paralogs, connected by undirected edges (e.g. panel (4) in Fig. 3A). The OrthNet module compare the open reading frame (ORF) length of a protein-coding node with the median value for neighboring ones and identify nodes with truncated ORFs (Fig. 3A). The OrthNet module uses Markov clustering (MCL) (van Dongen and Abreu-Goodger 2012), based on edge weights assigned according to edge properties, aiming to divide large networks that are often a result of expanded gene families with a large number of paralogs into smaller clusters likely derived from a single ancestral locus (Supplementary Text S2). Each cluster of orthologs, separated by MCL, is given an OrthNet ID and represented as an ortholog network or an OrthNet (Supplemtary Methods M3). Finally, the module can search with a user-defined pattern of ortholog copy numbers, edge characteristics, and network topology, to retrieve all OrthNets sharing a given set of evolutionary events (Supplementary Methods M4). Several selected examples of OrthNets representing different evolutionary histories are shown in Figure 3A.

**Figure 3.**
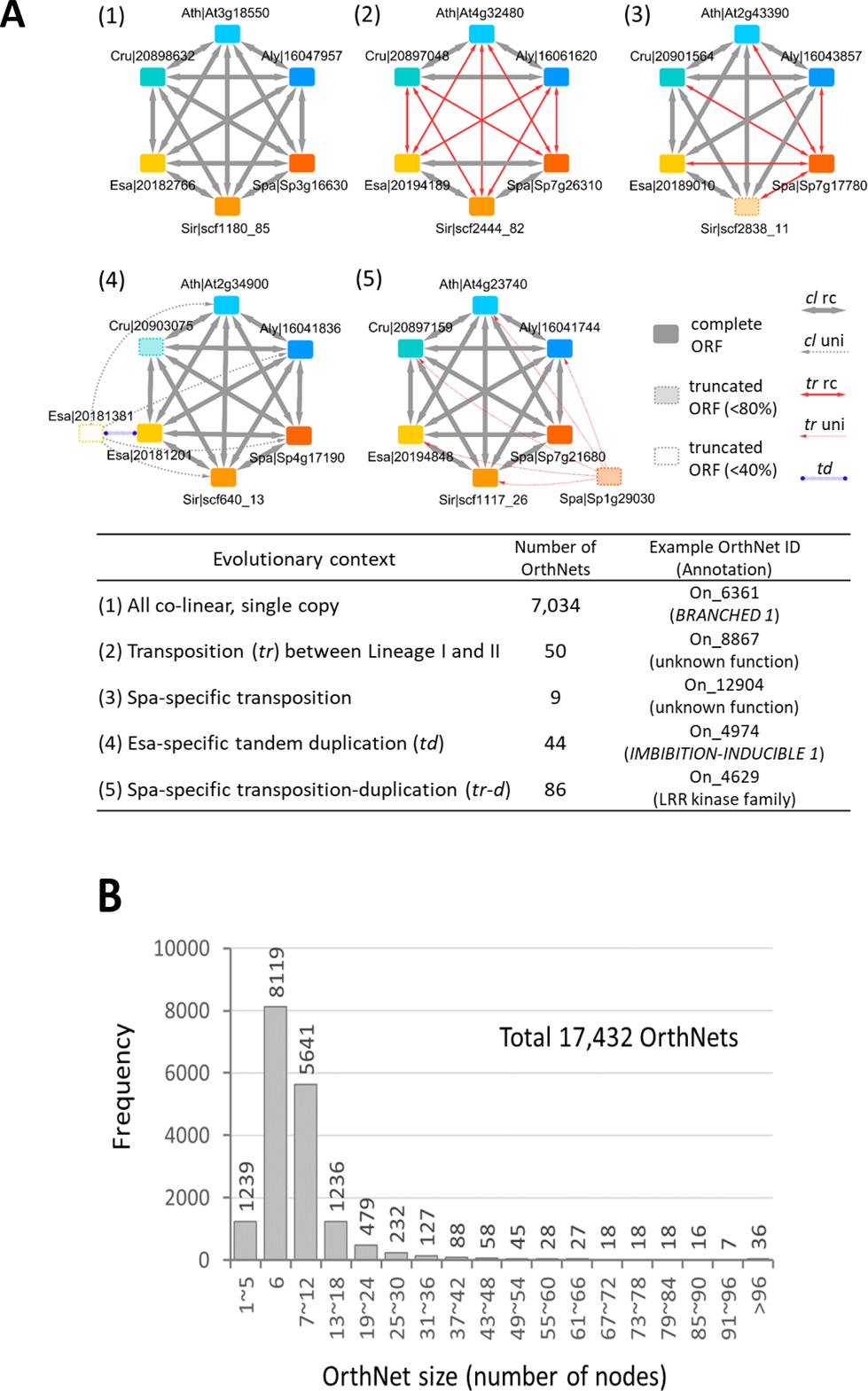
The OrthNet module encodes the evolutionary history of an orthologous gene group into a network. **(A)** OrthNet examples representing five different evolutionary histories. Nodes are color-coded according to the species. Transparent nodes with dashed borders indicate loci with truncated ORFs, i.e. ORF sizes smaller than either 80% or 50% compared to the median ORF size of nodes they are connected to. Edges show properties either co-linear (*cl*) or transposed (*tr*), reciprocally (rc) or uni-directionally (uni). Tandem duplicated (*td*) paralogs are connected by undirected edges. The lower panel shows the ID and annotation for representative OrthNets, as well as the number of OrthNets representing the same evolutionary history among 17,432 OrthNets identified for the six genomes. **(B)** A histogram showing the size distribution of OrthNets.

### Identification of OrthNets among six Brassicaceae genomes

We tested the CLfinder-OrthNet pipeline on the six Brassicaceae genomes using parameters and input files as described in Methods. The CLfinder module summarizes all reciprocal query-target genome pairwise analyses as exemplified for the six Brassicaceae genomes in Table 1. For simplicity, we considered *cl_end* loci pairs as *cl* in this summary. All query-target genome pairs showed a comparable number of *cl* loci pairs, ranging from 19,015 (Table 1, Sir-Aly) to 24,296 (Table 1, Aly-Ath). The number of *cl* pairs follows the division of the Lineage I (Table 1, Aly, Ath, and Cru) and II (Table 1, Esa, Sir, and Spa), with higher numbers observed between query-target pairs within each Lineage. The number of *tr* loci pairs was proportional to the repeat contents of the query genomes. For example, *A. lyrata* and *E. salsugineum* are the query genomes with the highest content of *tr* pairs (Table 1, Aly and Esa), which correlated with the higher content of repeats in these two genomes than in *A. thaliana, C. rubella*, or *S. parvula* genomes (“TE(%)” row in Table S1). When *S. irio* was the query, the proportion of *nd* pairs was higher than all other genomes (Table 1, Sir), because it had the most fragmented genome assembly among the six genomes. The entire CLfinder results for all query-target genome pairs is in Supplementary Dataset S1.

**Table 1.** Summary of CLfinder results showing pairwise comparisons among 6 crucifer species

| Query species | # protein-coding genes | CL type<sup>a</sup> | Target species |  |  |  |  |  | # *td*<sup>b</sup> events<br>(# *td* genes) |
| --- | --- | --- | --- | --- | --- | --- | --- | --- | --- |
|  |  |  | Aly | Ath | Cru | Esa | Sir | Spa |  |
| Aly | 32,657 | *cl* |  | 24,296 | 23,055 | 21,416 | 19,988 | 21,032 |  |
|  |  | *tr* |  | 4,881 | 5,375 | 6,668 | 8,104 | 6,478 | 2,163 |
|  |  | *ls* |  | 2,876 | 3,611 | 3,954 | 3,902 | 4,530 | (5,733) |
|  |  | *nd* |  | 604 | 616 | 619 | 663 | 617 |  |
| Ath | 27,206 | *cl* | 23,436 |  | 22,683 | 21,187 | 19,821 | 20,851 |  |
|  |  | *tr* | 2,431 |  | 2,804 | 4,032 | 5,355 | 4,064 | 1,747 |
|  |  | *ls* | 1,339 |  | 1,719 | 1,987 | 2,030 | 2,291 | (4,770) |
|  |  | *nd* | 0 |  | 0 | 0 | 0 | 0 |  |
| Cru | 26,521 | *cl* | 22,371 | 22,836 |  | 20,906 | 19,350 | 20,436 |  |
|  |  | *tr* | 3,036 | 2,836 |  | 4,338 | 5,817 | 4,267 | 1,752 |
|  |  | *ls* | 950 | 666 |  | 1,112 | 1,154 | 1,646 | (4,996) |
|  |  | *nd* | 164 | 183 |  | 165 | 200 | 172 |  |
| Esa | 26,351 | *cl* | 20,384 | 20,884 | 20,460 |  | 19,699 | 20,612 |  |
|  |  | *tr* | 4,465 | 4,137 | 4,460 |  | 5,431 | 4,046 | 1,646 |
|  |  | *ls* | 1,452 | 1,274 | 1,377 |  | 1,146 | 1,631 | (4,461) |
|  |  | *nd* | 50 | 56 | 54 |  | 75 | 62 |  |
| Sir | 32,524 | *cl* | 19,015 | 19,538 | 19,068 | 19,728 |  | 19,766 |  |
|  |  | *tr* | 3,062 | 2,860 | 2,998 | 2,722 |  | 2,697 | 1,795 |
|  |  | *ls* | 5,520 | 5,148 | 5,496 | 5,054 |  | 4,225 | (4,586) |
|  |  | *nd* | 4,927 | 4,978 | 4,962 | 5,020 |  | 5,836 |  |
| Spa | 26,847 | *cl* | 19,849 | 20,358 | 19,934 | 20,380 | 19,546 |  |  |
|  |  | *tr* | 2,830 | 2,452 | 2,718 | 2,534 | 4,097 |  | 1,242 |
|  |  | *ls* | 3,649 | 3,541 | 3,688 | 3,432 | 2,526 |  | (3,049) |
|  |  | *nd* | 519 | 496 | 507 | 501 | 678 |  |  |

<sup>a</sup> Co-linear (*cl*), transposed (*tr*), lineage-specific (*ls*), or not determined due to too small genome scaffold (*nd*)<sup>a</sup> Using CLfinder parameters {window\_size, num\_CL\_trshld, gap\_CL\_trshld} = { 20, 3, 20 }<sup>b</sup> Tandem duplication (*td*) detected using the parameter max\_TD\_loci\_dist = 4

The OrthNet module combined all pairwise CLfinder analyses and developed networks of orthologs, or OrthNets, representing the evolutionary history of each set of orthologous loci as the network topology (Fig. 3). For an analysis of N genomes, a perfect polygon (e.g. hexagon in the current study) with each of N nodes connected to other nodes by N-1 bidirectional solid gray edges represents a single-copy co-linear orthologous gene in all genomes (Fig. 3A, panel (1)). We identified a total of 7,034 OrthNets that showed single-copy loci co-linear to each other in all genomes. Panel (2) of Fig. 3A is an example from 50 OrthNets with co-linearity found within each of the Lineage I and II while loci between the two Lineages were transposed, representing a transposition event following the lineage divergence. Panel (3) shows one of the nine OrthNets with only the locus in *S. parvula* transposed compared to all other species. We found 44 OrthNets with the same evolutionary history depicted in panel (4), i.e. *E. salsugineum-specific* tandem duplication, and 86 OrthNets for *S. parvula-specific* transposition-duplication (*tr-d*) events shown in panel (5) of Fig. 3A. The OrthNet module also compares the ORF size of a node with the median ORF size of all other orthologous nodes to which the node is connected, to identify truncated ORFs (e.g. panels (3), (4), and (5) in Fig. 3A). We included all OrthNets together with the CLfinder results identified among the six Brassicaceae genomes in Supplementary Dataset S1.

An OrthNet may include a disproportionately large number of duplicated gene loci in specific genomes. For example, an OrthNet showing *A. lyrata-specific tr-d* events included 82 nodes representing additional *A. lyrata* transposed-duplicated paralog copies (Figure S4). Such duplication events, as well as large gene families where exact reciprocal ortholog pairs were hard to identify among multiple paralogs, may result in an OrthNet with a large number of nodes. However, more than 85% of OrthNets contain the same or less than 12 nodes per OrthNet (14,849 out of total 17,432 OrthNets), likely derived from single ancestral loci with duplications restricted in a subset of the six Brassicaceae genomes (Fig. 3B).

### Characterization of lineage-specific and shared *tr-d* events among Brassicaceae genomes

We used the “search OrthNet” functionality (Fig. S2, “search_OrthNet.py” and Supplementary Methods M4) to detect OrthNet representing either lineage-specific *tr* and *tr-d* events unique to each of the six Brassicaceae genomes, or *tr* and *tr-d* shared by more than one genome (Figure 4 and 5 and Dataset S2). The number of OrthNets that showed *tr* events was smaller than those with *tr-d* events for all subset of genomes including lineage-specific events (Table S1). This observation agrees with the postulation that a *tr* event is a result of a deletion of the original/donor copy after a *tr-d* event (Wicker et al. 2010).

In OrthNets including *tr-d* events, we identified the original donor or co-linear (CL) copy, or copies if the donor locus included tandem duplications, and the acceptor or transposed (Tr) copies, based on properties of edges connecting each of the duplicated paralogs to its neighboring nodes in other genomes (Figure 4A and 5A). Fig. 4A represents OrthNets with *S. parvula* and *C. rubella* lineage-specific *tr-d* events for orthologs of the *WRKY72* and *AGL87*, respectively. The CL copy (Fig. 4A, “CL copy”) was a part of the hexagon and mostly reciprocally co-linear to its ortholog nodes (Fig. 4A, “Orthologs”) from other genomes. A Tr copy was connected to ortholog nodes in the hexagon through uni-directional *tr* edges (Fig. 4A, “Tr copies”). An OrthNet may contain a single lineage-specific *tr-d* event as in the OrthNet for *WRKY72* (Fig. 4A, left) or multiple events featuring one CL copy associated with multiple Tr copies. Also, Tr copies may further undergo tandem duplication as shown in the OrthNet for *AGL87* (Fig. 4A, right).

**Figure 4.**
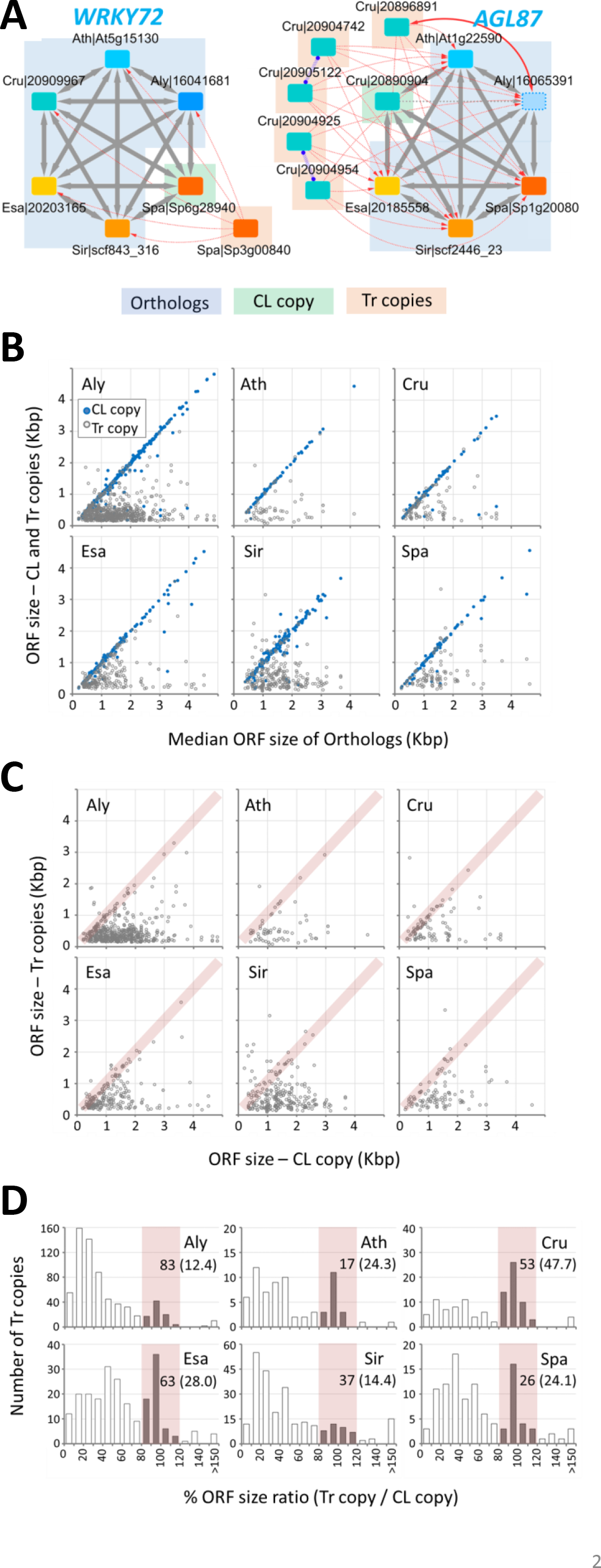
Characterization of lineage-specific transposition-duplication (*tr-d*) events among the six Brassicaceae genomes. **(A)** Examples of OrthNets with *tr-d* events specific for *S. parvula (WRKYDNA-BINDING PROTEIN 72/WRKY72*) and *C. rubella* (*AGAMOUS-LIKE 87/AGL87*). Within a *tr-d* event, OrthNet identifies the original donor copy (CL copy), which is co-linear to orthologs in other genomes (Orthologs), and the transposed and duplicated copies (Tr copies). Node and edges are as described in Fig. 3A. **(B)** ORF size comparison between all loci involved in a *tr-d* event (i.e. both CL and Tr copies) with the median of Orthologs. Blue dots indicate CL copies. **(C and D)** ORF size comparison between Tr copies and their corresponding CL copy within each of the *tr-d* events, as a scatterplot (C) and a histogram of ORF size ratio (D). Pink shades indicate Tr copies with complete ORFs whose sizes are comparable (±20% in proportion) to that of the CL copy. Panel D also shows numbers and percentages (in parentheses) of Tr copies with complete ORFs below the species labels. The entire list of OrthNets showing lineage-specific *tr-d*, including CL and Tr copies, is in Supplementary Dataset S2.

We compared the ORF sizes between CL and Tr copies with the median ORF size of the orthologs from other genomes in the hexagon for all OrthNets representing lineage-specific *tr-d* events. We observed a conservation of ORF sizes between most CL copies and their co-linear orthologs (Fig. 4B, blue dots), while the majority of Tr copies had truncated ORFs (Fig. 4B, gray dots). We also found a small proportion of Tr copies which had ORFs that were of similar size to their respective CL copy (Fig. 4C and 4D, pink rectangles). The distribution of the ORF size ratio between Tr and the CL copy showed peaks at 80∼120% (Fig. 4D, pink rectangles). These Tr copies that showed conservation in maintaining the original size of the ORFs were more abundant in *A. thaliana, C. rubella, E. salsugineum*, and *S. parvula.* These contributed to more than 24% of all Tr copies found in lineage-specific *tr-d* events in these genomes (Fig. 4D).

For *tr-d* events shared between any pair within the six genomes, the OrthNet module identified two categories with different evolutionary contexts: (1) parallel *tr-d* events independently occuring in two genomes (Figure 5, “Ind-parallel”) and (2) *tr-d* events where Tr copies from two genomes showing co-linearity between them (Fig. 5, “Tr-cl”). Fig. 5A depicts examples of OrthNets including *tr-d* events in the two categories. We found a total of seven and six OrthNets with *tr-d* events in “Ind-parallel” and “Tr-cl” categories, respectively, shared between *E. salsugineum* and *S. parvula.* Genomes with higher TE and repetitive sequence contents, such as *A. lyrata, S. irio*, and, to a lesser extent, *E. salsugineum*, included more “ind-parallel” *tr-d* events shared with other genomes (Fig. 5B, left panel). Among Tr copies in “ind-parallel” *tr-d* events, the proportion of complete ORFs (i.e. ORF size within ±20% of the ORF of the corresponding CL copy) were comparable to Tr copies in LS *tr-d* events (Fig. 5C, left panel and Fig. 4D).

**Figure 5.**
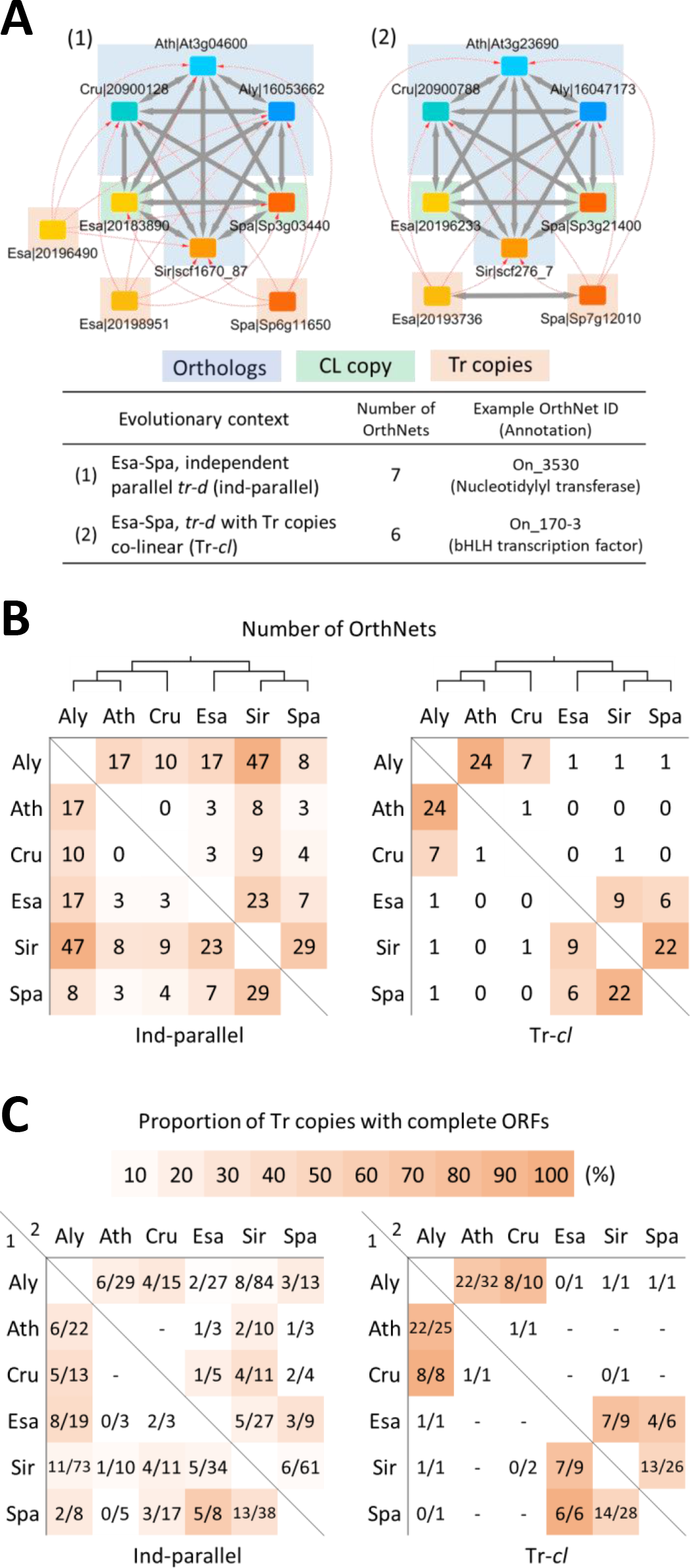
Transposition-duplication (*tr-d*) events shared by a pair of Brassicaceae genomes **(A)** Example OrthNets with *tr-d* events shared by *E. salsugineum* and *S. parvula*, representing two categories: (1) independent-parallel (“Ind-parallel”) *tr-d* events and (2) *tr-d* events with Tr copies colinear to each other (“Tr-cl”). Node and edges are as described in Fig. 3A. **(B)** Number of OrthNets shared by pairs of genomes in “Ind-parallel” and “Tr-cl” categories, with the cladogram of the six crucifer genomes on the top. Heatmap colors visualize the rank of each cell based on OrthNet numbers in each category. **(C)** Proportion of Tr copies with complete ORFs (i.e. ORF size ±20% of the CL copy in proportion) within OrthNets presented in (B). The genome 1 (row)-genome 2 (column) position shows the number of Tr copies with complete ORF / total Tr copies in genome 1, for all OrthNets with *tr-d* shared by genome 1 and 2. For example, the Aly-Ath and Ath-Aly positions in “Ind-parallel” category indicate 6 out of 29 *A. lyrata* (Aly) Tr copies and 6 out of 22 *A. thaliana* (Ath) Tr copies, respectively, have complete ORFs in the 17 “Ind-parallel” *tr-d* events shared by *A. lyrata* and *A. thaliana.* Heatmap colors indicate the percentage of Tr copies with complete ORFs, for cells with >5 total Tr copies.

The “Tr-cl” type *tr-d* events were mostly found between pairs of more recently diverged genomes, e.g. *A. lyrata-A. thaliana* and *S. irio-S. parvula.* The number of “Tr-cl” *tr-d* events detected between Lineage I and II genomes were very low (Fig. 5B, right panel). This observation was consistent with the notion that such a rare event must involve a *tr-d* event before the divergence of the two Lineages followed by deletions in all species except for the two genomes compared. The proportion of Tr copies that retained complete ORFs compared to the CL copy in “Tr-cl” type *tr-d* events was higher (≥50%) than that found for “ind-parallel” type *tr-d* events (Fig. 5C).

### Tr copies with complete ORFs were rare, but significantly more frequent than random chance

We hypothesized that selection has favored conservation of beneficial Tr copies to preserve the ORF in additional gene copies (Fig. 4C and D, pink rectangles; Fig 5C), while majority of the Tr copies were either originally duplicated incompletely or have undergone mutations over time that had led to truncated ORFs. An alternative hypothesis is that these Tr copies with complete ORFs may have been easier to duplicate in their complete form by random chance due to their smaller gene size. Indeed, genes associated with Tr copies with complete ORFs were significantly shorter than those with Tr copies that had truncated ORFs (Figures 6B and S5).

To test our hypotheses, we shuffled duplicated genomic regions and duplicated genes in *tr-d* events. Then, we compared the occurrences of randomized *tr-d* events showing complete duplication of the entire CL copy gene with those observed among actual *tr-d* events (Figure 6). First, to detect duplicated genomic regions in a *tr-d* event, we compared adjacent genomic regions, i.e. 5Kb up- and downstream regions, including the gene, of the CL copy and each of Tr copies. In this comparison, we searched for Homologous Genome Segments (HGSs) between the CL and Tr copy loci. As depicted in Fig. 6A, an incomplete *tr-d* event results in a HGS carrying only a part of the CL copy gene (HGS ⊅ CL copy), while in a complete *tr-d*, the HGS encompasses the entire CL copy gene (HGS ⊇ CL copy). Interestingly, we found a subset of complete *tr-d* events where the start and end positions of the HGS appeared to overlap with the start and end of the CL copy gene (HGS ≈ CL copy). We named this subset “gene-only” *tr-d* (Fig. 6A) since the sequence homology was not detectable in the intergenic region further from the CL copy coding regions by more than 20% of the CL copy gene size. A total of 224 complete *tr-d* events showed a shift towards longer HGSs, while their CL copy genes were significantly shorter (P<0.001, two-tailed t-test), compared to those in the 1,166 incomplete *tr-d* events (Fig. 6B).

**Figure 6.**
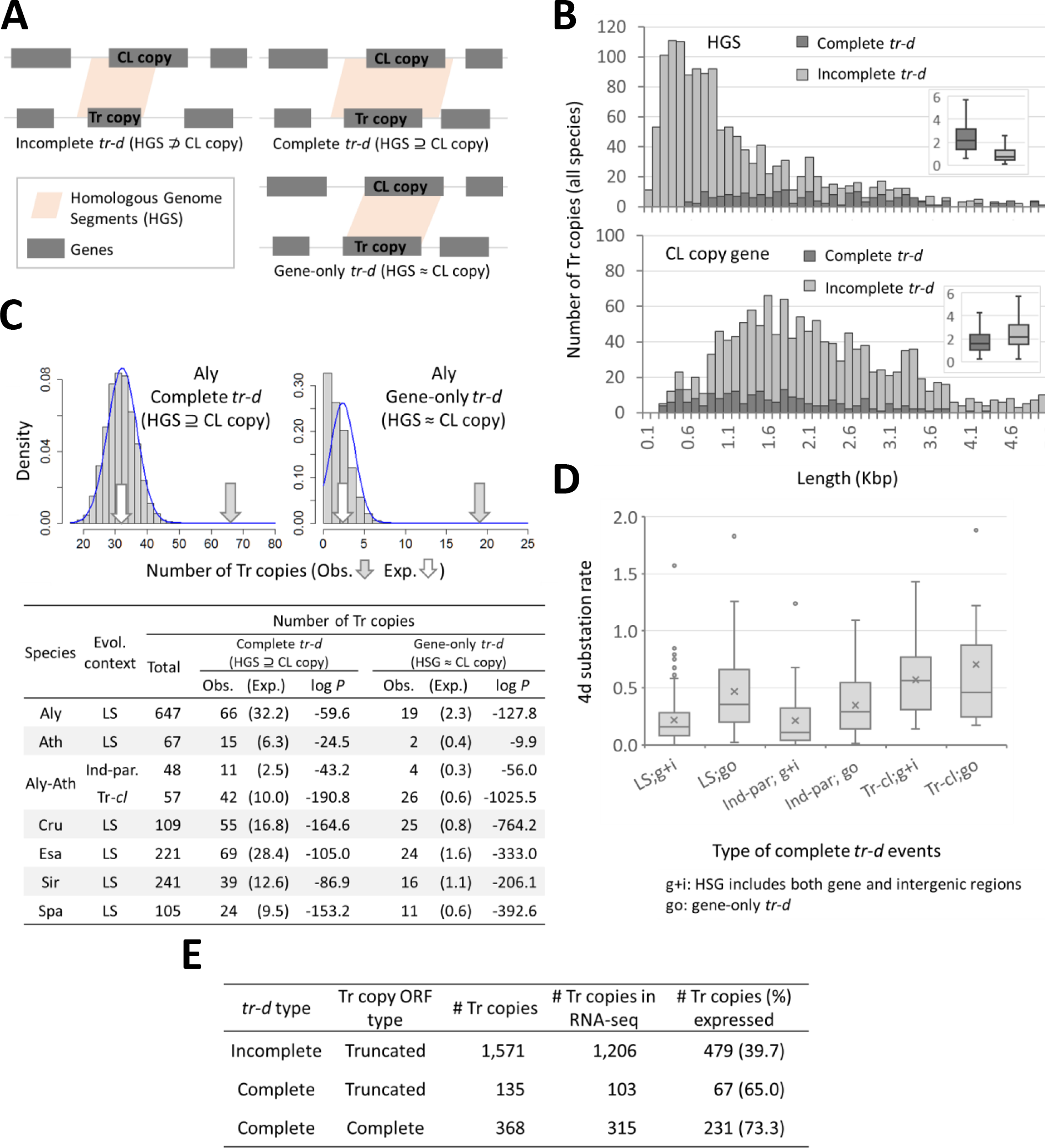
Characterization of duplicated genomic regions in transposition-duplication (*tr-d*) events. **(A)** We Identified Homologous Genome Segments (HGSs) between the CL and Tr copy genes and adjacent genomic regions (±5Kbp) in a *tr-d* event as described in Methods. A *tr-d* event is either complete or incomplete, based on whether the HGS included the full CL copy gene or not. A subset of complete *tr-d* events had HGS coinciding with the start and end of the CL copy gene without extending to the intergenic regions (“gene-only” *tr-d).* **(B)** Histograms and box-and-whisker plots (inlets) showing size distributions of HGSs and CL copy genes for complete and incomplete *tr-d* events. **(C)** Comparison of observed occurrences (Obs.) and expected occurrences (Exp.) of complete and gene-only *tr-d* events. Upper panel shows the distribution of expected occurrences from 10,000 random shuffling of HGSs and CL copy genes for *A. lyrata* (Aly)-specific complete and gene-only *tr-d* events. Fitting the random shuffling results to normal distributions (upper panel, blue curves) generated P-values of observed occurrences for *tr-d* events unique to each genome and the genus Arabidopsis (lower panel). **(D)** Four degenerate site (4d) substitution rates between ORFs of CL and Tr copy genes in different types of complete *tr-d* events. Complete *tr-d* events were either gene-only (“go”) or with HGSs detected in both gene and intergenic regions (“g+I”). We compared lineage-specific (LS) and shared *tr-d* events that are either independent-parallel (“Ind-par”) or with Tr copies co-linear to each other (“Tr-cl”). Lines and “x” marks in the box indicate medians and means, respectively. **(E)** Proportion of Tr copy genes with expression evidences (RNA-seq FPKM>0) in all *tr-d* events either lineage-specific or shared by a pair of genomes. The *tr-d* type is as described in (A) and Tr copy ORF type is as in Fig. 4C and D (pink shade) and Fig. 6C. *S. irio* genes were excluded due to the lack of RNA-seq data.

Following random shuffling of all HSGs and CL copy genes as described in Methods, we counted the occurrences of incomplete, complete, and gene-only *tr-d* events for each iteration. Fig. 6C shows the comparison between the observed and expected occurrences of complete and gene-only *tr-d* events, where expected values were the mean values from 10,000 iterations. Assuming a normal distribution for the expected values, we estimated the P-value for the observed numbers of complete and gene-only LS *tr-d* events for each genome. Both complete and gene-only *tr-d* events were much more frequent than expected due to random chance. The gene-only *tr-d* events had smaller p-values than complete *tr-d* events in all categories tested except in *A. thaliana* lineage-specific *tr-d* events (Fig. 5C, table in the lower panel). We observe a smaller number of lineage-specific *tr-d* events in *A. thaliana* than in any target genome. This may be a result of *A. thaliana* and *A. lyrata* being the closest among all pairs, included in the same genus. Hence, we included the *Arabidopsis* genus-specific *tr-d* events into consideration, which led to comparable numbers and enrichment of complete and gene-only *tr-d* events to other genomes (Fig. 5C, “Aly-Ath”).

Random occurrence of duplications cannot explain the observed proportion of complete *tr-d* events, which in >90% of the cases also resulted in complete ORFs in the Tr copy loci (e.g. Figs. 4C and 4D, pink shades). More likely, the observed proportion of complete and gene-only duplications was the sequential result of random duplications and selective retention of beneficial coding regions over time. This explanation is consistent with 4d substitution rates between complete ORFs of Tr copies and CL copies in *tr-d* events. Higher 4d substitution rates, as a proxy for older duplications, were found between ORFs of Tr and CL copy pairs in gene-only *tr-d* events (Fig. 6D, “go”). This was contrasting to CL copies in *tr-d* events where HGSs comprised both gene and intergenic regions (Fig. 6D, “g+i”), for both lineage-specific (Fig. 6D, “LS”) and indepedent parallel (Fig. 6D, “Ind-par”) shared *tr-d* events. The 4d substitution rates associated with “Tr-cl” type shared *tr-d* events (Fig. 6D, “Tr-cl”) showed median values comparable or higher than the median 4d substitution rates that represent the divergence between *A. thaliana* and Lineage II genomes (Fig. 2D). This further agreed with the notion that a “Tr-cl” type *tr-d* event was derived from duplications dated prior to the divergence of genomes that shared the events.

Complete *tr-d* events also included a higher number of Tr copy genes that showed evidence of expression compared to incomplete *tr-d* events (Fig. 6E). Incomplete *tr-d* was associated with most of the Tr copies with truncated ORFs, which comprised the majority of Tr copies in both lineage-specific *tr-d* (Fig. 4C and 4D) and independent parallel *tr-d* events shared by a pair of genomes (Fig. 5C). Out of total 1,706 Tr copies with truncated ORFs, only 135 were derived from complete *tr-d* events, in which the ORFs were most likely truncated by null mutations after the duplication (Fig. 6E). We found no enrichment of single exon genes, a signature of retrotransposons, among *tr-d* events (Figure S6).

### Genes associated with lineage-specific and shared *tr-d* events

Table S2 presents a partial list of OrthNets associated with lineage-specific *tr-d* events for each of the six Brassicaceae genomes, selected based on the most number of Tr copies with complete ORFs and expression evidences, except for *S. irio*, for which RNAseq data was not available. For each OrthNet listed in Table S2, we included numbers of Tr copies tandem duplicated, with complete ORFs, and with expression evidences, as detailed in Supplementary Text S3. The complete list of OrthNets including lineage-specific *tr-d* events is available in Dataset S2. We described genes and gene ontology terms enriched among them in lineage-specific *tr-d* events in Supplementary Text S3 and Dataset S3.

We selected the largest OrthNets with *E. salsugineum-specific tr-d* events (Table S2) and independently visualized the extent of gene duplications using the GEvo tool in the CoGE database (Lyons and Freeling 2008) (Figure 7). The *E. salsugineum* genome included six copies of *SALT TOLERANCE 32 (SAT32*), which exists as a single copy in each of the other Brassicaceae genomes. Among five Tr copies detected for *EsSAT32*, we found three tandem duplicates (Fig. 7A, “Tr copies”). Four of the Tr copies had complete ORFs and three of them showed expression in either root or shoot tissues (Fig. 7A and Table S2). The GEvo plot illustrates extensive sequence similarity among all loci and adjacent genomic regions that are reciprocally co-linear among them (Fig. 7B, *AtSAT32, SpSAT32*, and *EsSAT32;1*). Similar patterns were observed in comparisons with the *S. irio, C. rubella*, and *A. lyrata* co-linear orthologs (data not shown). The *EsSAT32* Tr copies (Fig. 7B, *EsSAT32;2/3/4/5*) represented examples of gene-only *tr-d* events (Fig. 6A), where sequence similarities were restricted to the expected border regions of the gene models (i.e. ±20% of the coding region size). Interestingly, the Tr copies *EsSAT32;3/4/5* also exhibited intron loss, resulting in 9, 4, and 1 exons, respectively, compared to the 13 exons in the CL copy *EsSAT32;1* (Fig. 6B), while maintaining high deduced amino acid similarities over most of the coding region (Figure S7). Among all *EsSAT* paralogs, the highest average expression was observed for one of the Tr copies, *EsSAT32;2* (Supplementary Dataset S1, OrthNet ID ON_2516 and gene ID 20186362).

**Figure 7.**
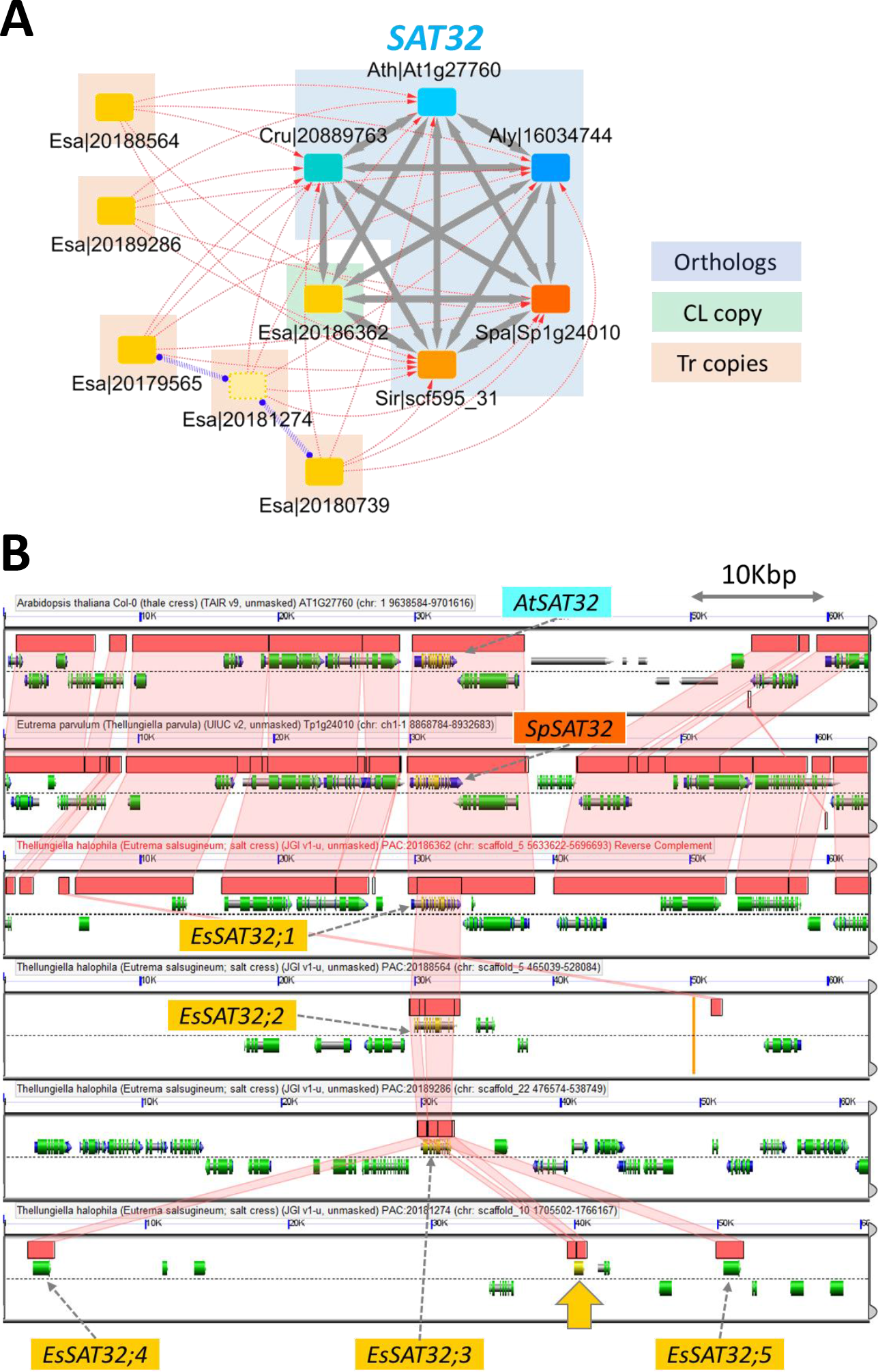
Examples of lineage-specific *tr-d* events. **(A)** OrthNet for the *SALT-TOLERANCEE 32* (*SAT32*). Nodes and edges are as described in Fig. 3A. The OrthNet showed *E. salsugineum-specific tr-d*, with three of the five Tr copies tandem duplicated. (B) Comparison of *SAT32* loci and adjacent ±30Kbp genomic regions between *A. thaliana, S. parvula*, and *E. salsugineum* as a GEvo plot (https://genomevolution.org/r/maxx). Pink shades connect Homologous Genomic Segments (HGSs) between genomes, while gene models, mRNAs, and coding sequences are depicted in gray, blue, and green, respectively (for detailed GEvo legends, see https://genomevolution.org/wiki/index.php/GEvo). *EsSAT32;1* is the CL copy (Esa|20186362), while *EsSAT32;2/3/4/5* indicate the four Tr copies with complete ORFs. The yellow arrow marks the position of Esa|20181274, which contains a truncated ORF. *EsSAT32;3/4/5* showed intron losses without compromising gene products (see text and Figure S7 for details).

For *tr-d* events shared by multiple genomes, we present the entire list of such OrthNets for all pairs of genomes, as well as those associated with Lineage I and II-specific *tr-d* events, in Supplementary Dataset S2. The *tr-d* events shared by *E. salsugineum* and *S. parvula* were of particular interest, because they may indicate signatures of convergent evolution between these two species independently adapted to high salinity (Dittami and Tonon 2012; Oh et al. 2012). Table S3 lists all OrthNets with “Ind-parallel” and “Tr-cl” type *tr-d* events (as defined in Fig. 5A) shared between these two extremophytes. We also included all Lineage II (i.e. *E. salsugineum, S. irio*, and *S.* parvula)-specific *tr-d* events that had truncated Tr copy ORFs only for *S. irio* (Table S3, marked by superscript “e”). Among the “Ind-parallel” tr-d events detected, three out of seven events were associated with stress signaling or response-related functions (Table S3, *CDPK1, 5PTASE11*, and *ABI1*). However, none of them included complete Tr copy ORFs in both *E. salsugineum* and *S. parvula.* Interestingly, the “Tr-cl” category included more loci with Lineage II-specific *tr-d* followed by truncation of the Tr copy ORF in *S. irio*, leaving complete ORFs in the Tr copy loci only for *E. salsugineum* and *S. parvula.* Here, we found loci encoding orthologs of a putative basic helix-loop-helix (bHLH) type transcription factor, a NAC transcription factor (NAC058), and a calcineurin B-like protein 10 (CBL10). All Tr copies encoding these regulatory proteins showed expression evidence in both halophytes (Table S3).

## DISCUSSION

### A systematic identification of ortholog loci with the same evolutionary history

In this study, we developed the CLfinder-OrthNet pipeline in an attempt to systematically identify all gene loci among multiple genomes that underwent the same set of evolutionary events, such as gene duplications and transpositions in a certain lineage or multiple lineages that are either mono-, para-, or polyphyletic.

Previous works have suggested network representation of synteny among orthologs as an effective method to combine and summarize synteny blocks identified by all-to-all pairwise comparisons among multiple genomes (Zhao and Schranz 2017). Synteny networks connecting co-linear orthologs from multiple genomes with undirected edges traced the evolutionary path of a gene family (Zhao et al. 2017). This approach has been used to compare the extent of gene duplications and lineage-specific expansion of gene families between mammalian and plant genomes (Zhao and Schranz 2018). While the CLfinder module similarly performs all-to-all pairwise analyses to detect co-linearity in gene order, OrthNets detected by the CLfinder-OrthNet pipeline are different from synteny networks (Zhao and Schranz 2017) in a number of ways. For example, while synteny networks connected co-linear nodes with undirected edges, OrthNets connected nodes with directional edges with co-linearity or lack of it (i.e. transposed) encoded as edge properties. An OrthNet includes orthologs connected by reciprocal edges as well as paralogs derived from duplications connected by uni-directional edges to their neighboring nodes found in other genomes (Fig. 3A, panel (4) and (5); Fig. 4A and 5A). We aimed to separate each OrthNet into a unit that represents a group of orthologs and paralogs likely derived from a single ancestral locus, by employing Markov clustering (MCL) (Supplementary Text S2). We chose MCL to control edge weights to prefer undirected tandem duplicated edges and reciprocal edges over uni-directional edges during the clustering process (Supplementary Methods M3). In this way, each of the majority of OrthNets, e.g. >85% of all OrthNets in case of the six Brassicaceae genomes (Fig. 3B), represents the evolutionary history of genes derived from a single ancestral locus as the network topology. Essentially, OrthNets enable detection of all loci from multiple genomes that share the same evolutionary history by a search using a given network topology as the query (e.g. Fig. 3A and 5A). We used this functionality to characterized transposition-duplication (*tr-d*) events in six Brassicaceae genomes, as a proof-of-concept (Supplementary Methods M4).

### Transposition-duplication as a major mechanism for erosion of co-linearity

For the transposition and transposition-duplication (*tr-d*) of non-TE gene loci, two types of models, retrotransposon-associated and DNA repair or replication-associated models have been suggested as the main mechanisms (Hastings et al. 2009; Robberecht et al. 2013). A *tr-d* event derived from retrotransposons often leads to duplication of single exon genes (Morgante et al. 2005; Cusack and Wolfe 2007; Abdelkarim et al. 2017). Transposition-duplication may also arise during the non-homologous end-joining (NHEJ) repair process of DNA double-strand breaks (DSB), where a short sequence motif may act as an anchor to a foreign sequence to fill-in a gap (Wicker et al. 2010). In agreement with this model, a previous comparison of *A. lyrata* and *A. thaliana* found a significant enrichment of flanking repeats, as short as 15bps, among transposed genes (Woodhouse et al. 2010). The correlation between the proportion of query gene loci showing distal displacement (Fig. S1C, *d_n, n+1_>20* or “Diff Chr”) and overall TE contents of the query genome, rather than divergence time (Fig. S1D), supports the DSB-repair model. Higher TE contents likely provide a higher frequency of short repeat anchors required for the NHEJ DSB-repair, and TE activities themselves may also cause the DSB that lead to such repairs (Wicker et al. 2010). The DSB-repair model can explain *tr-d* of multi-exon genes, which constitute the majority of *tr-d* events found in the current study (Fig. S6).

In lineage-specific *tr-d* events retrieved by OrthNet, we found a subset of transposed-duplicated gene loci (Fig. 4A, Tr copies) retaining similar ORF sizes compared to their respective donor locus (Fig. 4A, CL copy), as well as to orthologs in other species (Fig. 4B-D). The DSB repair model of *tr-d* suggests that the duplicated region may start and end virtually at any random position in a genome, given that the short sequence motif needed for the repair is likely ubiquitously available and can be as short as several nucleotides (Wicker et al. 2010; Woodhouse et al. 2010). However, our simulation reveled that both “complete” and “gene-only” *tr-d* events were far more frequent than what was expected from a random duplication model alone (Fig. 6C). We are not aware of a gene duplication mechanism that preferably duplicates non-TE, protein-coding, multi-exon genes as entire units. Rather, our observation common to all six tested crucifer genomes is likely a result of random *tr-d* events (e.g. through DSB repair), followed by accumulation of mutations throughout the duplicated regions, except where the complete coding sequences were selectively retained. Supporting this notion, Tr copy genes with complete ORFs were more frequent among shared *tr-d* of older “Tr-cl” type events (Fig. 5C and 6D). These were also more likely to be expressed, hence less likely to be pseudogenes, compared to Tr copies with truncated ORFs (Fig. 6E). See Supplementary Text for further discussions on the age of “gene-only” *tr-d* events (Supplementary Text S4) and on *tr-d* frequencies and TE contents (Supplementary Text S5). Overall, our analyses depicted the landscape of *tr-d* events among Brassicaceae genomes, where the majority of *tr-d* were incomplete, while small numbers of *tr-d* including complete Tr copy ORFs and gene-only *tr-d* were likely to have resulted from random duplication events followed by selective retention of coding sequences over time.

### Search for extremophyte-specific *tr-d* events using OrthNets

One possible application of the CLfinder-OrthNet pipeline is to retrieve orthologs sharing evolutionary events unique to a lineage with a specific trait or multiple lineages exhibiting a convergent trait, e.g. the two extremophyte *S. parvula* and *E. salsugineum.* As detailed in Supplementary Text S6, these two genomes have been identified with gene copy number and structural variations compared to *A. thaliana* that were associated with stress-adapted traits (Oh et al. 2009; Oh et al. 2010; Ali et al. 2012). In this study, we used the CLfinder-OrthNet pipeline to identify 63, 26, and 14 ortholog groups showing gene copy number increases through *tr-d* events specific to *E. salsugineum, S. parvula*, and both, respectively (Fig. 4D and Table S3). These numbers are orders of magnitude fewer than previous searches from a pairwise comparison to *A. thaliana* (Oh et al. 2014), signifying the vastly improved resolution in finding extremophyte-specific events.

The OrthNet for the *SALT-TOLERANCE 32 (SAT32*) locus (Figure 7A and Table S2, ON_2516) represents the largest *E. salsugineum-specific tr-d* event. *SAT32* encodes a transcription regulator, whose expression level positively correlated with the survival rate of the model plant *A. thaliana* under salt stress (Park et al. 2009). Three of the four *EsSAT32* paralogs with complete ORFs exhibited intron losses (Figs. 7B and S7). Intron losses and smaller transcript sizes are reported to enable regulation of expression timing in Drosophila and mouse (Hao and Baltimore 2013; Jiang et al. 2014). It is not clear whether “gene-only” *tr-d* events (Fig. 7B) among *EsSAT32* paralogs is indicative of reverse transcriptase-mediated duplication leading to intron losses (William Roy and Gilbert 2006) or different rate of mutation between gene and intergenic regions. Either way, such variation in intergenic regions including promoter regions may lead to sub-functionalization (Wang, Wang, et al. 2012). At least three *EsSAT32* paralogs exhibited different basal expression strengths in root and shoot tissues (data not shown).

A notable example of *S. parvula-specific tr-d*, with copy number increases of complete ORFs, is the *ZRT/IRT-LIKE PROTEIN 3 (ZIP3*) locus encoding a zinc transporter (Table S2). This particular *tr-d* may be a signature of an adaptation in *S. parvula*, to soils that are highly saline and also depleted in micronutrients such as zinc and iron in central Anatolia (Cakmak et al. 1999; Eide 2005). See Supplementary Text S7 and S8 for discussions on genes involved in *tr-d* unique to each extremophyte, as well as *tr-d* shared by the two extremophytes.

### Concluding remarks: CLfinder-OrthNet, a flexible toolkit for comparative genomics

The CLfinder-OrthNet pipeline, in a proof-of-concept application, successfully encodes more than 85% of entire loci among six Brassicaceae genomes into OrthNet units in which evolutionary histories of genes derived from single ancestral loci can be traced (Fig. 3B). Using a network topology-based search (Supplementary Method M4), we identified groups of orthologs, represented as OrthNets that share the same evolutionary histories (Fig. 3A), including *tr-d* unique to any subset of the six Brassicaceae genomes (Fig. 4 and 5, Supplementary Dataset S2).

As detailed in Supplementary Text S9, CLfinder-OrthNet offers multiple options to apply the pipeline flexibly depending on target genomes and goals of the study. The sensitivity and stringency of co-linearity detection are adjustable by controlling parameters depending on the range of target genomes. The CLfinder module can use results from any method of sequence clustering and comparison, as well as genomic features other than protein-coding genes, as inputs. Moreover, the two modules can be used separately. For example, researchers can use the CLfinder module to quickly summarize the distribution of co-linear, tandem duplicated, and transposed genes among multiple genomes (e.g., Table 1), while the OrthNet module can accept locus-level synteny information from other methods to generate OrthNets.

Overall, the CLfinder-OrthNet pipeline offers a flexible toolkit to compare the arrangement of gene and other genomic features among multiple genomes. Future applications include, but not limited to, tracing evolutionary histories of a gene or gene families; inference of orthology based on both sequence homology and co-linearity; studying incongruence between sequence homology and synteny; (Supplementary Text S2), and identification of candidate copy number variations associated with certain traits (e.g., as discussed in Supplementary Text S6-S8).

## MATERIALS AND METHODS

### Genome and gene models

We obtained genome annotations of *A. lyrata* (Aly, version 1.0), *A. thaliana* (Ath, v. “TAIR10”), *C. rubella* (Cru, v. 1.0), and *E. salsugineum* (Esa, v. 1.0) from Phytozome v. 11 (http://genome.jgi.doe.gov/), while genomes of *S. irio* (Sir, v. 0.2; CoGE genome id 19579) and *S. parvula* (v. 2.0) were downloaded from CoGE (https://genomevolution.org/coge/) and thellungiella.org (http://thellungiella.org/data/), respectively. For *S. irio* annotation, we used a combination of RepeatMasker, a BLASTN search versus Repbase v. 20170127 (http://www.girinst.org/repbase/) and OrthoMCL (Li et al. 2003) to further filter out lineage-specific clusters of genes encoding components of transposons. To generate a species tree of crucifer genomes including the six target species, we used Agalma (Dunn et al. 2013) which built a maximum likelihood tree based on 14,614 alignments of homologous protein-coding gene clusters. Each cluster contained sequences from more than four crucifer genomes. We analyzed co-linearity erosion among crucifer genomes as described in Supplementary Methods M1.

### CLfinder-OrthNet on Brassicaceae genomes

The three inputs, parameters, and detailed CLfinder process to identify co-linearity among best-hit pairs is described in Supplementary Methods M2. For the analysis of six crucifer genomes, CLfinder parameters window_size (*W*) = 20, num_CL_trshld (*N*) = 3, gap_CL_trshld (*G*) = 20, and max_TD_loci_dist (*T*) = 4 were decided based on the distribution of protein-coding gene locus content in the scaffolds of the most fragmented genome (S. *irio*) and the results from the analysis of co-linearity erosion. The CLfinder process (Fig. S3) was performed for all possible query-target pairs for the six crucifer species using a wrapper script in the CLfinder module (Fig. S2, “CLfinder_multi.py”).

The OrthNet module connected all best-hit ortholog and, if any, tandem duplicated paralogs in a locus with co-linearity information as the edge property. The resulting networks were further divided into smaller unit, using MCL with default edge weights as detailed in Supplementary Methods M3. The final network units (OrthNets) were converted to .sif file format for visualization using Cytoscape (http://cytoscape.org/).

### Analysis of transposition-duplication (*tr-d*) events

OrthNets including a *tr-d* events uniquely found in subsets of the six crucifer genomes were identified using the “search-by-seeds” functionality of the OrthNet module, as described in Supplementary Methods. Within an OrthNet with a *tr-d* event, the *tr-d* donor or “CL copy” was the node connected to orthologous nodes with the most *cl* edges, while the remaining were *tr-d* acceptors or “Tr copies.” When multiple CL copies existed due to tandem duplication, we used the one with the longest ORF as the representing CL copy. Homologous Genome Segments (HGSs) were detected between the gene and ±5Kb intergenic regions of the CL copy and each of Tr copies, using LASTZ with chaining as previously described (Oh et al. 2014). A *tr-d* event was “complete” if the HGS included the entire CL copy gene. A “gene-only” *tr-d* was defined as a complete *tr-d* event with the size of HGS less than 120% of the CL copy gene. We determined the expected occurrences of complete and gene-only *tr-d* by random shuffling and overlapping of HGSs and CL copy genes. The distribution of such occurrences from 10,000 iterations was fitted to a normal distribution to calculate the P-value of the observed occurrence, using the fitdistr function in R MASS package (https://cran.r-project.org/web/packages/MASS).

The 4d substation rates were calculated for all CL and Tr copy pairs where the Tr copy contined a complete ORF, using codeml (Yang 2007) and a custom script (Fig. S2, “pairwiseKs_by_codeml.py”). To determine Tr copies with expression evidence, we used RNA-seq data for leaf and root tissues obtained from studies by Wang et al. (2016) for *A. lyrata, A. thaliana*, and *C. rubella*, and Oh et al. (2014) for *A. thaliana* and *S. parvula* as well as samples prepared for the current study (for *E. salsugineum*, BioProject ID PRJNA63667) essentially as previously described (Oh et al. 2014). FPKM values of representative transcript models were estimated using Stringtie (v. 1.3.1c) with the ‘-e’ option (Pertea et al. 2016), after RNAseq reads were aligned to the genome using HISAT2 (v. 2.0.5) (Pertea et al. 2016).

### Availability of software

All custom scripts and the CLfinder-OrthNet pipeline are available on GitHub (https://github.com/ohdongha/CL_finder).

